# Acid and inflammatory sensitisation of naked mole-rat colonic afferent nerves

**DOI:** 10.1101/636571

**Authors:** James R.F. Hockley, Katie H. Barker, Toni S. Taylor, Gerard Callejo, Zoe M. Husson, David C. Bulmer, Ewan St. J. Smith

## Abstract

Acid sensing in the gastrointestinal tract is required for gut homeostasis and the detection of tissue acidosis caused by ischaemia, inflammation and infection. In the colorectum, activation of colonic afferents by low pH contributes to visceral hypersensitivity and abdominal pain in human disease including during inflammatory bowel disease. The naked mole-rat (*Heterocephalus glaber*; NMR) shows no pain-related behaviour to subcutaneous acid injection and cutaneous afferents are insensitive to acid, an adaptation thought to be a consequence of the subterranean, likely hypercapnic, environment in which it lives. As such we sought to investigate NMR interoception within the gastrointestinal tract and how this differed from the mouse (*Mus Musculus*). Here we show the presence of calcitonin gene regulated peptide (CGRP) expressing extrinsic nerve fibres innervating both mesenteric blood vessels and the myenteric plexi of the smooth muscle layers of the NMR colorectum. Using *ex vivo* colonic-nerve electrophysiological recordings we show differential sensitivity of NMR, compared to mouse, colonic afferents to acid and the prototypic inflammatory mediator bradykinin, but not direct mechanical stimuli. In NMR, but not mouse, we observed mechanical hypersensitivity to acid, whilst both species sensitised to bradykinin. Collectively, these findings suggest that NMR colonic afferents are capable of detecting acidic stimuli, however, their intracellular coupling to downstream molecular effectors of neuronal excitability and mechanotransduction likely differs between species.

## Introduction

The gastrointestinal (GI) tract coordinates the digestion of food, absorption of nutrients and evacuation of waste with acidification of the stomach contents a critical component of this process. Through compartmentalisation, sensory surveillance and specialised mucosal defence mechanisms, not only is the breakdown of food and elimination of ingested pathogens achieved through acidification in the foregut, but also the delicate gut microbiota-host symbiosis of the hindgut maintained. It is clear that when gastric acid regulation is lost then significant pathogenesis can occur, including acid-related diseases such as gastro-eosophageal reflux disease, gastroduodenal ulceration, dyspepsia and gastritis [1].

Sensory neurones innervating the GI tract are central to the feedback regulation of gastric acid secretion and can additionally detect tissue acidosis caused by inflammation, ischaemia and microbial activity [2,3], often resulting in visceral hypersensitivity and abdominal pain [4]. Whilst luminal pH varies along the length of the healthy human gut (with lower pH found in the stomach and colon [5,6]), both surgical intervention and disease (e.g. chronic pancreatitis and inflammatory bowel disease [5,7,8]) can also result in abnormal acidification of the gut.

The naked mole-rat (*Heterocephalus glaber*; NMR) has adaptations enabling it to not only survive but prosper in the subterranean (thus likely hypercapnic and hypoxic) environment in which it lives, including profound hypercapnia and hypoxia resistance[9,10]. Many of these adaptations have led to altered sensory processing of external stimuli, for example the NMR shows no pain-related behaviour to subcutaneous injection of acid and capsaicin [11], lacks an itch response to histamine [12] and shows no thermal hyperalgesia in response to a variety of stimuli, including nerve growth factor [11,13]. These adaptations are believed to provide a fitness advantage to living in a subterranean environment, for example, the likely high CO_2_ environment of NMR nests would evoke noxious stimulation of C-fibres through acidosis in almost any other rodent [14]. Whilst clearly valuable in supporting its lifestyle in its niche, detection of acid by sensory neurones innervating the gut is required for maintenance of gut homeostasis and for the detection of tissue acidosis. In other rodent species, viscerally-projecting afferent fibres can sense tissue acidosis by specialised ion channels [1] including acid-sensing ion channels (ASICs; which respond to mild acidification), transient receptor potential vanilloid subtype (TRPV1; which are gated by severe acidosis), ionotropic purinoceptor ion channels (e.g. P2X_3_) and two-pore domain potassium channels (e.g. TASK, TRESK, TREK and TRAAK subtypes); additionally, a number of proton-sensing G protein-coupled receptors exist that are also sensitive to mild acidification (e.g. Gpr68, Gpr4, Gpr132 and Gpr65)[15]. Recent single-cell RNA-sequencing analysis of colonic sensory neurones shows discrete expression of such acid-sensitive ion channels with differing populations of mouse colonic afferents, suggesting functional specialism [16]. Compared to mice, NMRs display a similar expression profile of ASICs throughout the nervous system [17], and of those analysed, with the exception of ASIC3, NMR acid sensors show similar activation profiles to those of mice [13,18,19]. NMR TRPV1 is also expressed in sensory afferents and shows similar proton sensitivity to mouse TRPV1 [18]. The NMR acid-insensitivity is likely due to an amino acid variation in the voltage-gated sodium channel 1.7 subunit (Na_V_1.7), which results in acid anaesthetising, rather than activating their cutaneous sensory neurones [18]. Considering the unusual cutaneous acid-insensitivity of the NMR, it is of interest to determine how GI sensory surveillance and detection of visceral tissue acidosis occurs in this species, especially considering the growing reputation of the NMR as a model of healthy ageing [20] and the perturbation of GI function that occurs with ageing [21]. In order to investigate this, we examined the sensory innervation of the NMR colon and made electrophysiological recordings in both NMR and mouse from the lumbar splanchnic nerve innervating the colorectum, and applied noxious mechanical and chemical stimuli, including acid and bradykinin, a prototypic inflammatory mediator.

## Materials and Methods

### Animals

Experiments were performed in C57BL6/J mice (6-41 wks; 3F, 6M) and subordinate NMR (25-198 wks; 2F, 7M). Mice were conventionally housed with nesting material and a red plastic shelter in temperature-controlled rooms (21 °C) with a 12 h light/dark cycle and access to food and water *ad libitum*. NMRs were bred in-house and maintained in an inter-connected network of cages in a humidified (~55 %) temperature-controlled room (28 °C) with red lighting (08:00-16:00) and had access to food *ad libitum*. In addition, a heat cable provided extra warmth under 2-3 cages/colony. NMRs used in this study came from two different colonies. Experiments were conducted under the Animals (Scientific Procedures) Act 1986 Amendment Regulations 2012 under Project Licenses (70/7705 & P7EBFC1B1) granted to E. St. J. Smith by the Home Office and approved by the University of Cambridge Animal Welfare Ethical Review Body.

### Immunohistochemistry

The colorectum was dissected free before opening along the mesenteric border and pinning flat in a Sylgard lined dissection tray. After fixing in Zamboni’s fixative (2 % paraformaldehyde / 15 % picric acid in 0.1 M phosphate buffer; pH 7.4) overnight, the mesentery and mucosa were dissected free from the muscle layers. Small 1.5 cm x 1.5 cm sections were subsequently washed in 100 % DMSO (3 x 10 min) and phosphate buffered saline (PBS; 3 x 10 min). Tissues were blocked with antibody diluent (10 % donkey serum, 1 % bovine serum albumin (BSA) in 0.2 % Triton X-100) for 1 h, then primary antibodies were applied overnight at 4 °C. The following day, tissues were washed (PBS, 3 x 10 min), donkey anti-rabbit IgG-AF488 (1:500, Life Technologies A21206) antibody applied for 2 h, washed (PBS; 3 x 10 min), mounted and coverslipped. Primary antibodies used were rabbit anti-calcitonin gene-related peptide (1:5000, Sigma C8198) and rabbit anti-protein gene product 9.5 (1:500, Abcam ab10404). No labelling was observed in control sections where primary antibody was excluded. Tissues were imaged using a Leica SP5 confocal microscope and z-stack reconstructions of nerve fibres within different layers of the NMR gut produced with ImageJ (v1.51a, NIH).

### Electrophysiology recordings of visceral afferent activity

Colonic nerves innervating the colorectum of mouse and NMR were isolated and electrophysiological activity was recorded as previously described [22]. Mice were humanely killed by cervical dislocation of the neck and cessation of circulation. NMRs were humanely killed by CO_2_ exposure followed by decapitation. For both species, the colorectum with associated lumbar splanchnic nerve was dissected free from the animal and transferred to a recording chamber superfused with carbogenated Krebs buffer (in mM: 124 NaCl, 4.8 KCl, 1.3 NaH_2_PO_4_, 2.5 CaCl_2_, 1.2 MgSO_4_.7H_2_O, 11.1 glucose and 25 NaHCO_3_; 7 ml/min; 32-34 °C). The colorectum was cannulated and perfused with Krebs buffer (100 μl/min) enabling distension of the colon by closure of the out-flow. The Krebs buffer was supplemented with nifedipine (10 μM) and atropine (10 μM) to inhibit smooth muscle activity and with indomethacin (3 μM) to restrict endogenous prostanoid production. Multi-unit electrophysiological activity of the lumbar splanchnic nerve rostral to the inferior mesenteric ganglia was recorded using a borosilicate glass suction electrode. Signals were amplified and bandpass filtered (gain 5K; 100-1300 Hz; Neurolog, Digitimer Ltd, UK) and digitised at 20 kHz (micro1401; Cambridge Electronic Design, UK) before display on a PC using Spike 2 software. The signal was digitally filtered online for 50 Hz noise (Humbug, Quest Scientific, Canada) and action potential firing counts were determined using a threshold of twice the background noise (typically 100 μV).

### Electrophysiological protocols

Tissues were stabilised for 30 min before noxious intraluminal distension pressures were applied by blocking the luminal out-flow of the cannulated mouse or NMR colorectum. The pressures reached are above threshold for all known visceral afferent mechanoreceptors [23] and evoke pain behaviours in rodents *in vivo* [24]. Mechanosensitivity and chemosensitivity were investigated using a combined sequential protocol. As such, a slow ramp distension (0-80 mmHg, 4-5 min) and set of 6 rapid phasic distensions (0-80 mmHg, 60 s at 9 min intervals) were applied as previously described [25] prior to bath superfusion of pH 4.0 Krebs buffer (50 mL volume) and a set of 3 phasic distensions (0-80 mmHg, 60 s at 9 min) to test for acid-induced acute mechanical hypersensitivity. After a 20 min wash-out period, 1 μM bradykinin was applied by bath superfusion (20 mL volume) and a further set of 3 phasic distensions were performed. Phasic distension protocols were automated using an Octaflow II perfusion system (ALA Scientific, USA) to standardise duration and intervals.

### Data analysis

Peak changes in firing rates of electrophysiological nerve recordings were determined by subtracting baseline firing (3 min before distension or drug application) from increases in nerve activity following distension or chemical stimuli. Statistical analysis was performed using two-way analysis of variance (ANOVA) followed by Holm-Sidak’s post hoc test in Prism 6 (GraphPad Inc., USA). Statistical significance was set at *P* < 0.05. Data are displayed as means ± SEM.

### Drugs

Stock concentrations of bradykinin (10 mM; water), nifedipine (100 mM; DMSO), atropine (100 mM; ethanol) and indomethacin (30 mM; DMSO) were dissolved as described, diluted to working concentration in Krebs buffer on the day of experiment as described above and were all purchased from Sigma-Aldrich.

## Results

### Gastrointestinal neuroanatomy of the NMR

We first compared the gross anatomy of the NMR and mouse GI tract. As the NMR is greatly long lived compared to the mouse, with a life expectancy of >30-years, we chose animals from both species that could be deemed adults (see Methods). Compared to the mouse, NMR GI length (from pyloric sphincter to anus) was significantly shorter in length (mouse, 37.8 ± 0.4 mm; NMR, 25.8 ± 1.2 mm, *P* = < 0.001, *N* = 3).

We next confirmed the presence of extrinsic sensory fibres innervating different layers of the NMR colorectum using immunohistochemistry. Equivalent staining in the mouse are widely available within the literature and we did not seek to duplicate them here [26]. Using antibodies raised against calcitonin gene-related peptide (CGRP) and protein gene product 9.5 (PGP9.5) we stained for neuronal fibres within flat-sheet whole-mount preparations of multiple layers of the NMR colon. Specifically, CGRP-positive extrinsic neuronal varicosities were identified encircling and tracking with blood vessels within the mesentery supplying the distal colon of NMR; such fibres likely contribute to the larger lumbar splanchnic nerve upon which these coalesce (Fig. 1B). Although NMR lack CGRP in cutaneous afferent neurones, this finding is in line with the observation that mesenteric arteries in NMR and the common mole-rat (*Cryptomus hottentotus*) express CGRP [27]. Neuronal fibres staining for PGP9.5 were also observed within the mesentery of NMR, again localised around blood vessels (Fig. 1E). The mucosa and submucosa were separated from the muscle (circular and longitudinal) layers. CGRP-positive, presumably extrinsic, sensory fibres were observed coursing through the myenteric plexi between these muscle layers (Fig. 1C). PGP9.5 staining revealed the myenteric soma and additional neuronal fibres within this layer of the NMR colon (Fig. 1F). Whilst PGP9.5-positive fibres were observed encircling the base of colonic villi (see Fig. 1G insert), CGRP labelling of these fibres was not seen (Fig. 1D). Interdigitating fibres within both the circular and longitudinal muscle layers are positive for PGP9.5 (Fig. 1H and 1I), with what is likely submucosal ganglia retained on the circular muscle layer after separation of the mucosa from this layer (Fig. 1H).

**Figure 1.**
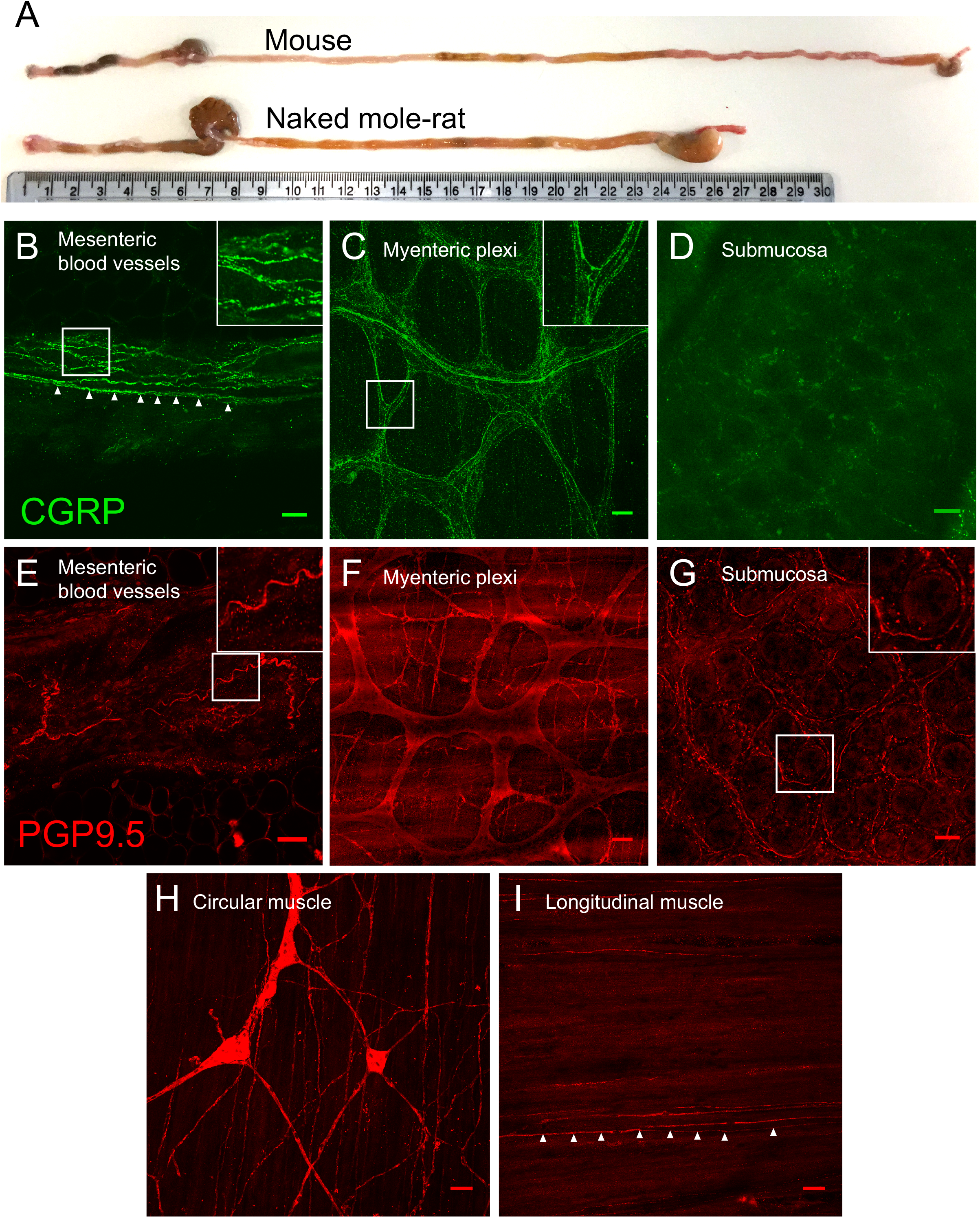
Extrinsic sensory innervation of NMR colorectum. A. Comparison of mouse and NMR gastrointestinal tracts from anus (*left*) to oesophagus (*right*), with a 30 cm ruler providing scale. Wholemount immunostaining for CGRP in the mesentery (B; *insert*, nerve fibres encircling a mesenteric blood vessel. *Arrows*, example nerve fibre on the blood vessel margin), myenteric plexi (C; *insert*, extrinsic nerve fibres infiltrating myenteric ganglia) and submucosa (D) of NMR. Equivalent nerve fibre staining was also observed with PGP9.5 in the mesentery (E; *insert*, nerve fibre surrounding mesenteric blood vessel), myenteric plexi (F), submucosa (G; *insert*, nerve fibre surrounding the base of a mucosal villi), circular muscle (H) and longitudinal muscle (I; *arrows*, nerve fibre innervating longitudinal muscle). Scale bar in each panel, 50 μm.

### Colonic afferent mechanosensitivity does not differ in the NMR compared to mouse

In order to understand whether the peripheral terminals of sensory neurones innervating the GI tract of the NMR possessed altered acid and inflammatory sensitivity compared to mouse, we made *ex vivo* multi-unit electrophysiological recordings of lumbar splanchnic nerve activity using a suction electrode from the colorectum of both NMR and mouse. The lumbar splanchnic nerve innervates the colorectum and is a pathway through which pain is the predominant conscious sensation transduced [28]. The colorectum, once dissected free from the animal, was cannulated and both luminally perfused and bath superfused with Krebs buffer, thus allowing mechanical distension of the bowel or application of chemical stimuli, respectively.

We first investigated mechanosensitivity of visceral afferents in the NMR compared to mouse GI tract. There were no significant differences in the baseline spontaneous activity measured between NMR and mouse (3 min average: 7.2 ± 2.6 spikes/s vs. 8.3 ± 1.2 spikes/s, respectively, *N* = 9, *P* = 0.72, unpaired t-test). We applied known innocuous and noxious mechanical stimuli, firstly by way of a ramp distension (0 to 80 mmHg) and using repeat phasic distension (Fig. 2A). By using a slow ramp distension, we were able to assess visceral afferent responses across a range of physiologically-relevant distension pressures typically exposed to the rodent gut [29,30]. We observed no difference in the nerve firing recorded during ramp distension in NMR (e.g. at 80 mmHg, 27.9 ± 5.6 spikes/s) compared to mouse (e.g. at 80 mmHg, 31.2 ± 5.2 spikes/s; Fig. 2B, *P* = 0.51, *N* = 8-9, mixed-model ANOVA). We next applied repeat phasic distension of the colon to noxious (0-80 mmHg) pressures. As previously reported in mouse, we observed a rapid increase in nerve activity to initial phasic distension (82.3 ± 9.9 spikes/s) and significant adaptation during the 60 s distension (Fig. 2C; [22,25]). Following subsequent repeat distensions at 9 min intervals, tachyphylaxis occurred with a decrease in peak firing of 22.0 % by the sixth distension compared to the first. In NMR, afferent discharge reached an equivalent peak firing compared to mouse during the first distension and the degree of desensitisation during subsequent distensions was similar (22.2 % by the sixth distension, *P* = 0.98, *N* = 9, two-way repeated-measures ANOVA; Fig. 2C). Similar afferent responses to mechanical stimuli in NMR compared to mouse suggest that there is no intrinsic difference in the way sensory nerves transduce physiological and noxious mechanical stimuli.

**Figure 2.**
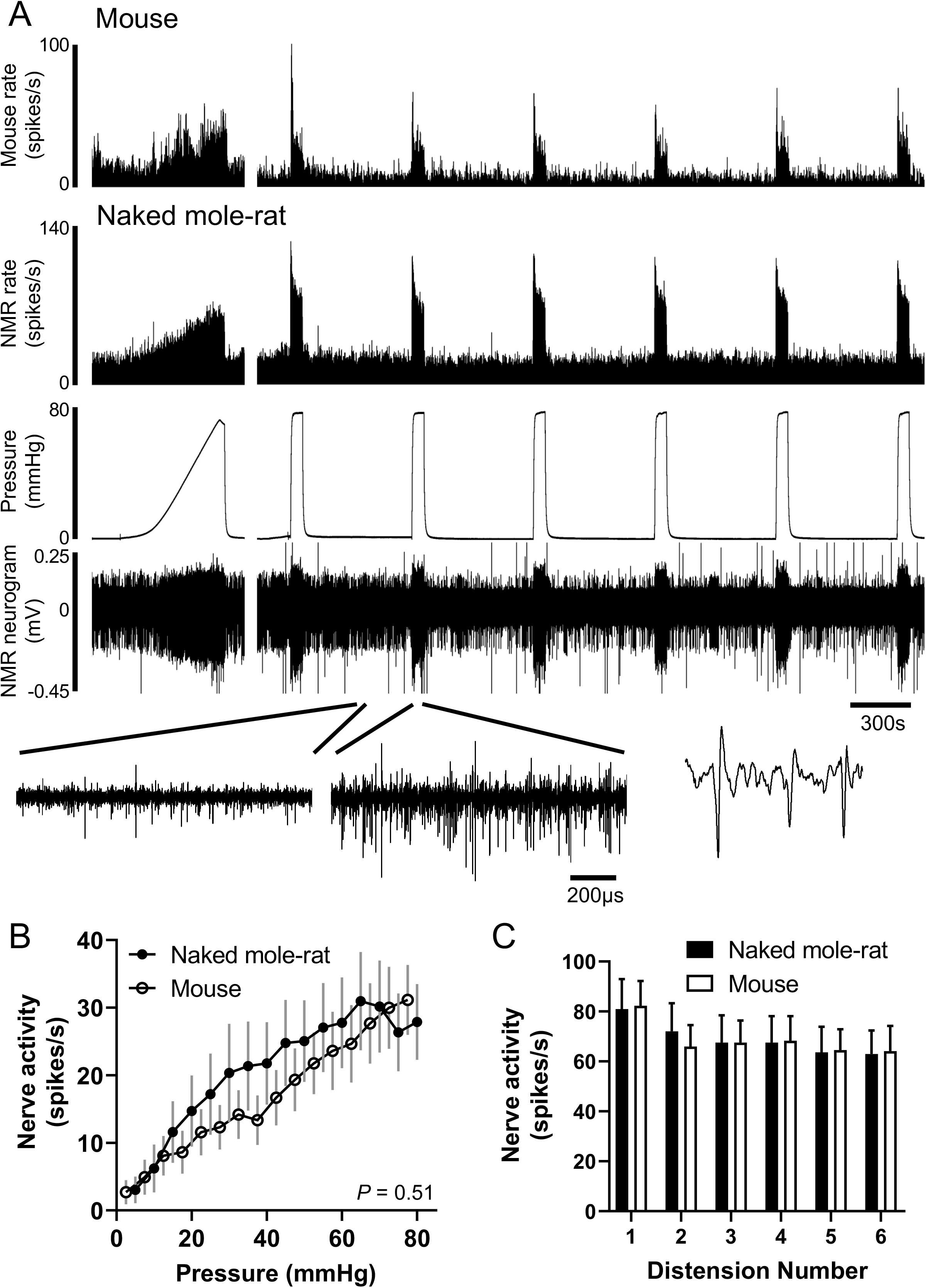
Colonic afferent responses to noxious ramp and repeat phasic distension in mouse and NMR. A. Example rate histograms of colonic lumbar splanchnic nerve activity from mouse and NMR with intraluminal pressure trace and neurogram trace following ramp distension (0 to 80 mmHg) and repeat phasic distension (0-80 mmHg, 60 s, 9 min intervals). Below, expanded neurogram traces showing NMR before and after phasic distension and an example trace showing three action potentials. B. Mean firing rates to ramp distension at 5 mmHg increments in mouse and NMR (*P* = 0.51, *N* = 8-9, mixed-model ANOVA). C. Average change in peak firing rate during repeat phasic distension in mouse and NMR (*P* = 0.98, two-way repeated-measures ANOVA).

### Extracellular acid evokes mechanical hypersensitivity in NMR but not mouse

Next, we investigated the effect of extracellular acid on visceral afferent firing and mechanical hypersensitivity to phasic distension (Fig. 3A). We chose a pH 4.0 stimulus to broadly activate acid-sensitive ion channels [15,31] and a stimulus that is capable of evoking pain both in humans and rodents when injected subcutaneously [11,32]. The vast majority of colonic sensory neurones possess inward sustained currents in response to low pH [33]. Bath superfusion of pH 4.0 to mouse colon directly excited visceral afferents evoking a peak firing increase of 30.0 ± 5.1 spikes/s returning to baseline firing rates after 1735 ± 60 s. Direct excitation of NMR visceral afferents as a result of acid was significantly lower compared to mouse, but return to baseline did not differ between species (12.5 ± 4.3 spikes/s and duration of 1545 ± 197 s, *P* = 0.02 and *P* = 0.37, respectively, *N* = 9, unpaired t-test; Fig. 3B). Immediately after returning to baseline, a set of three phasic distensions (60s, at 9 min intervals) was applied to test whether extracellular acid induced mechanical sensitisation. In agreement with previous studies in mouse, application of acid did not alter firing rates in response to any of the three subsequent phasic distensions when compared to the response prior to acid application (Fig. 3C; [34]). By contrast, extracellular acid caused significant mechanical sensitisation in the NMR, such that the response to phasic distension immediately after acid application was 48.3 % greater than before (*P* < 0.001, *N* = 9, 2-way ANOVA with Holm-Sidak’s post hoc test). This mechanical sensitisation was lost by the second post-acid phasic distention and by the third phasic distension afferent firing had recovered to baseline levels and was comparable to mouse (Fig. 3C). That low pH conditions, such as that observed during inflammation, can evoke robust mechanical hypersensitivity in NMR, but not mouse, suggests fundamental differences in the mechanism by which acid-sensitive receptors are coupled to mechanotransducers in the peripheral terminals of colonic sensory neurones.

**Figure 3.**
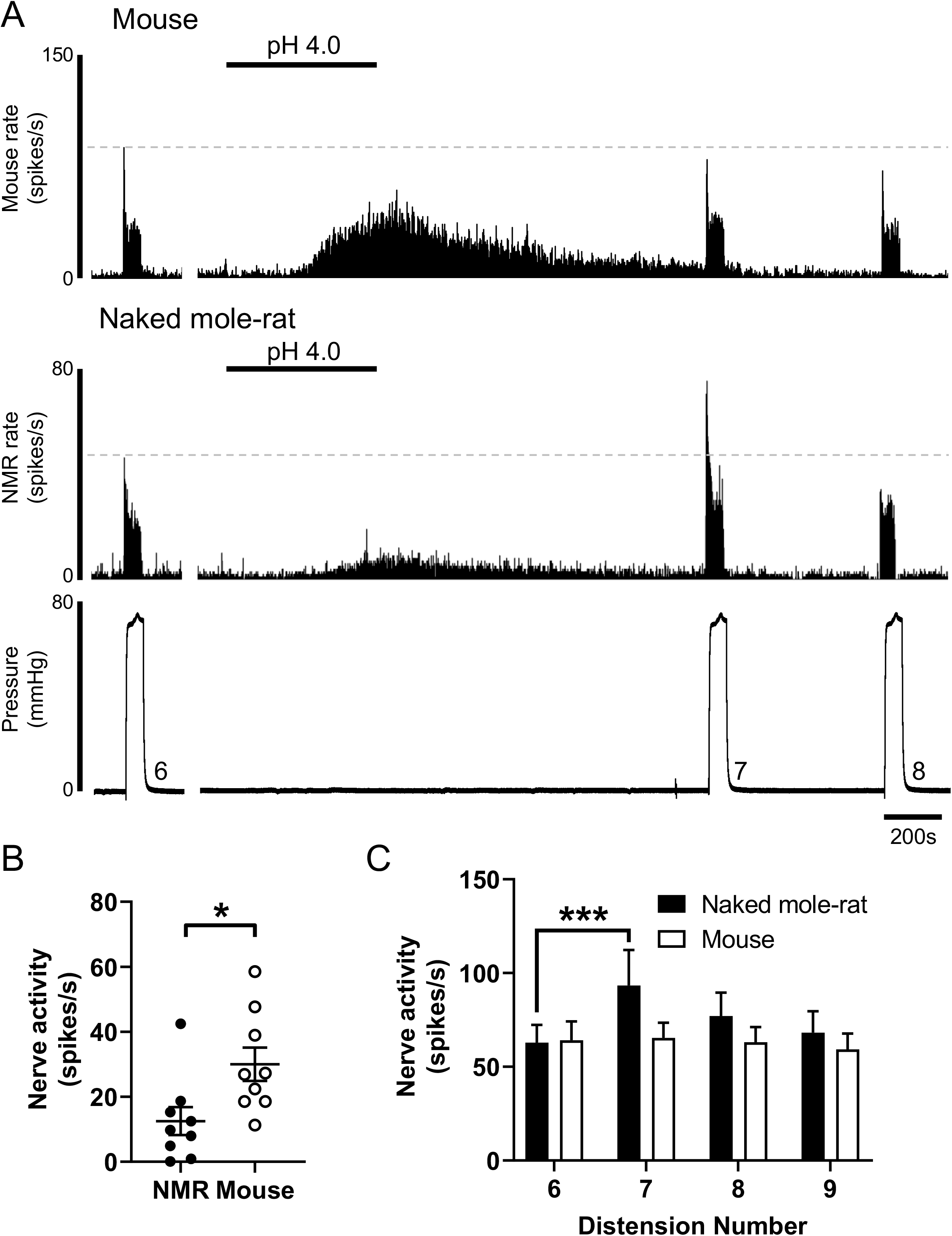
Extracellular acid evokes mechanical hypersensitivity in NMR, but not mouse. A. Example rate histograms of colonic splanchnic nerve activity from mouse and NMR with accompanying pressure trace showing bath superfusion of pH 4.0 Krebs buffer (50 mL) and subsequent repeat (x 3) phasic distension. B. Mean increase in peak firing after application of pH 4.0 (\**P* = 0.02, *N* = 9, unpaired t-test). C. Peak firing change to phasic distension after superfusion with pH 4.0 solution. The response to phasic distension in NMR, but not mouse, was significantly sensitised by acid (\*\*\**P* < 0.01, *N* = 9, two-way repeated-measures ANOVA with Holm-Sidak’s post-hoc).

### Afferent excitation to bradykinin is blunted in NMR, but mechanical sensitisation is unaffected

Given that inflammatory pain responses in NMR are blunted to some inflammatory stimuli [11], we investigated the ability for the prototypical inflammatory mediator, bradykinin, to not only activate, but evoke mechanical hypersensitivity in NMR visceral afferent fibres. Application of bradykinin (1 μM) by bath superfusion to mouse colonic afferents led to an increase in peak firing of 24.2 ± 3.9 spikes/s in agreement with previous studies in mouse and human colonic tissues (Fig. 4A; [35,36]). In NMR this was not the case, with peak firing only increased by 4.0 ± 1.5 spikes/s following addition of bradykinin (*P* < 0.001, *N* = 9, unpaired t-test; Fig. 4B). However, in both mouse and NMR, a robust mechanical hypersensitivity to phasic distension was observed immediately after bradykinin application, such that the response to 80 mmHg phasic distension was potentiated by 21 % in NMR and 29% in mouse (Fig. 4C).

**Figure 4.**
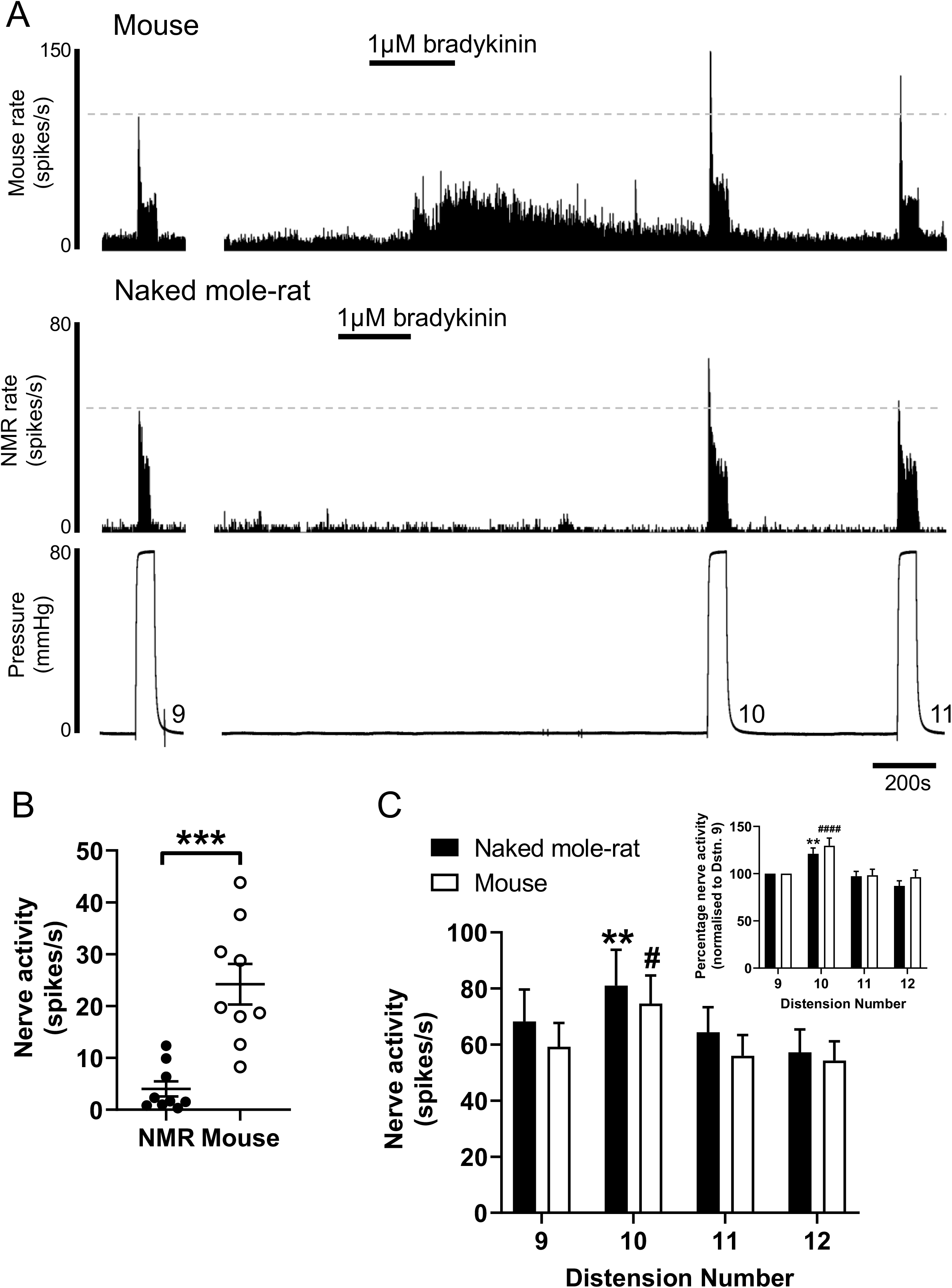
Colonic afferent excitation to bradykinin is blunted in NMR, but mechanical sensitisation is unaffected. A. Example rate histograms of colonic splanchnic nerve activity from mouse and NMR with accompanying pressure trace showing addition of 1 μM bradykinin (20 mL) and subsequent repeat (x 3) phasic distension. B. Mean increase in peak firing after application of 1 μM bradykinin (\*\*\**P* < 0.001, *N* = 9, unpaired t-test). C. Peak firing change to phasic distension after superfusion with 1 μM bradykinin. The response to phasic distension in both NMR and mouse was significantly sensitised by application of bradykinin (\*\**P* < 0.01 vs. 9^th^ distension in NMR, #*P* < 0.05 vs. 9^th^ distension in mouse, *N* = 9, two-way repeated-measures ANOVA with Holm-Sidak’s post-hoc). *Insert*, phasic distension responses in both NMR and mouse normalised to the prebradykinin distension response (\*\**P* < 0.01 vs. normalised 9^th^ distension in NMR, ####*P* < 0.0001 vs. normalised 9^th^ distension in mouse, *N* = 9, two-way repeated-measures ANOVA with Holm-Sidak’s post-hoc).

## Discussion

Acid sensing in the GI tract is necessary to maintain gut homeostasis by providing feedback for gastric and intestinal acid regulation, and secondly for detecting tissue acidosis caused by inflammation, infection and ischaemia during disease. Here we assessed the mechanical and chemical sensitivity of NMR and mouse colonic afferents in response to differing noxious stimuli.

In NMRs, the absence of thermal hypersensitivity induced by capsaicin and lack of histamine-induced scratching are thought to be due to a lack of cutaneous neuropeptides, such that both behaviours can be “rescued” by intrathecal administration of SP [11,12]. We show here by immunohistochemistry that CGRP is expressed within nerve fibres found encapsulating both blood vessels of the NMR colonic mesentery and myenteric plexi within the smooth muscle layers of the colon wall, which aligns with previous findings of CGRP-positive fibres innervating NMR mesenteric arteries [27]. By contrast, PGP9.5 staining identified nerve fibres within the submucosa that did not express CGRP, highlighting potential restricted penetration of extrinsic sensory fibres innervating the NMR colorectum. Such differences in sensory innervation termini did not manifest as altered mechanosensitivity to distension during baseline conditions of NMR colonic afferents compared to mouse. This suggests that visceral mechanotransduction is not significantly altered in the NMR.

We did observe greatly differing responses to both, the direct exposure of extracellular acid and to induced mechanical hypersensitivity, implicating altered integration of acid sensors, mechanotransducers and modulators of spontaneous afferent firing. NMR acid sensing differs significantly to other mammals. For example, in hippocampal and cortex neurones, the peak current density of NMR ASIC-like responses is reduced compared to mouse brain neurones [37]. In the peripheral nervous system, subcutaneous injection of acid (pH 3.5), capsaicin or histamine does not cause the nocifensive or pruriceptive behaviours in NMR that such stimuli characteristically induce in mice [11,12]. This acid insensitivity is a function of altered ASIC responses compared to mouse [19] and a variation in NMR Na_V_1.7, which renders the channel hypersensitive to proton-mediated block and therefore prevents acid-driven action potential initiation from the skin [18]. Such intrinsic differences in the sensitivity of NMR to acid may explain our observation of significantly lower firing rates in response to application of acid to NMR colonic afferents compared mouse. We have shown previously that pharmacological inhibition or genetic ablation of Na_V_1.7 in mouse does not impair colonic afferent firing or alter pain behaviours [25]. Therefore, if Na_V_1.7 is redundant in colonic afferents compared to those innervating the hindpaw, then it would be predicted that NMR colonic afferents would not be as insensitive to acid as their somatic equivalents. Whilst this hypothesis does not hold true for the direct action of acid on NMR colonic afferent firing, i.e. it is diminished compared to the mouse, it does fit with the lack of mechanical hypersensitivity observed following acid application in the mouse, which has been reported previously [34]. In contrast, the robust sensitisation observed in NMR colonic afferents to acid suggests differential coupling of molecular acid sensors to mechanotransducers compared to the mouse. Further studies would be required to elucidate the intracellular signalling cascades involved.

Responses to the inflammatory mediator bradykinin failed to activate NMR colonic afferents, but could induce a robust mechanical hypersensitisation comparable to the effects observed in mouse. Although we do not confirm bradykinin B2 receptor expression in NMR colonic afferents in this study, such mechanical hypersensitivity suggests that the B2 receptor activity is unimpaired. The bradykinin B2 receptor is a G_αs_ protein-coupled receptor which upon activation modulates the function of a number of molecular transducers (including K_V_7 [38], TRPV1 [39], TRPA1 [40], Ca^2+^-activated Cl^-^ channels [38,41] and K_Ca_ [42,43]) via downstream signalling cascades including increased intracellular calcium and PKC dependent phosphorylation. Differences in the activity or coupling to of these molecular transducers in the NMR may explain the altered response profiles compared to mouse.

Given our experimental paradigm, we believe our observations are most likely the result of species differences in molecular sensors and transducers responsible for action potential firing at the level of the primary afferent terminal, however we cannot exclude a number of confounding factors that may influence colonic afferent sensitivity. Firstly, whilst every effort was made to match the relative age range and sex of the NMR and mice used in this study, we acknowledge that age may have an impact on afferent sensitivity. For example, reduced responses of afferent firing to some stimuli are observed in tissues from older mice [44] and humans [45,46]. We accept that this may have introduced variability into our study, but given the relatively comparable spread in age of the individuals used from both species, we do not believe that this is responsible for the dramatic differences observed between species. Secondly, all NMR used in this study were subordinates and therefore their sexual maturity will have been suppressed. Sex steroid hormones including oestrogen can modulate visceral sensory function [47]. Without a direct comparison to breeding NMRs, which is a challenge due to the eusociality of the species, we are unable to exclude that the effects observed are not due to sexual immaturity. Thirdly, all electrophysiological recordings were made at 32-34 °C and as such represents a 2-3 °C difference from optimum body temperature for both species, albeit in opposing directions. Temperature affects afferent firing, however, the equivalence of responses to ramp and phasic distension acts as a positive control and suggests that NMR colonic afferents are capable of producing significant action potential firing in these conditions.

In summary, understanding how noxious pH is sensed and GI homeostasis is maintained in the NMR may help to inform our understanding of other model species and ultimately, GI acid sensing during human disease.

## Acknowledgements

The authors declare no competing financial interests. This work was supported by Rosetrees Postdoctoral Grant (A1296; JRFH and EStJS), BBSRC grant (BB/R006210/1; JRFH and EStJS), Versus Arthritis Pain Challenge Grant (RG21973; GC and EStJS), AstraZeneca PhD studentshipt (KHB), EMBO Long-Term Fellowship (ALTF1565-2015; ZH) and University of Cambridge Vice Chancellor’s Award (TST).

## Author Contributions

JRFH designed the research studies, conducted the experiments, acquired and analysed the data and wrote the manuscript. KHB, TST, GC and ZH acquired and analysed the data. DCB wrote the manuscript. EStJS designed the research studies and wrote the manuscript. All authors approve the final version of the manuscript.

